# A common stay-on-goal mechanism in anterior cingulate cortex for information and effort choices

**DOI:** 10.1101/2024.10.23.619920

**Authors:** Valeria V. González, Melissa Malvaez, Alex Yeghikian, Sydney Wissing, Melissa Sharpe, Kate M. Wassum, Alicia Izquierdo

## Abstract

Humans and non-humans alike often make choices to gain information, even when the information cannot be used to change the outcome. Prior research has shown the anterior cingulate cortex (ACC) is important for evaluating options involving reward-predictive information. Here we studied the role of ACC in information choices using optical inhibition to evaluate the contribution of this region during specific epochs of decision making. Rats could choose between an uninformative option followed by a cue that predicted reward 50% of the time vs. a fully informative option that signaled outcomes with certainty, but was rewarded only 20% of the time. Reward seeking during the informative S+ cue decreased following ACC inhibition, indicating a causal contribution of this region in supporting reward expectation to a cue signaling reward with certainty. Separately in a positive control experiment and in support of a known role for this region in sustaining high-effort behavior for preferred rewards, we observed reduced lever presses and lower breakpoints in effort choices following ACC inhibition. The lack of changes in reward latencies in both types of decisions indicate the motivational value of rewards remained intact, revealing instead a common role for ACC in maintaining persistence toward certain and valuable rewards.

**Graphical Summary:** 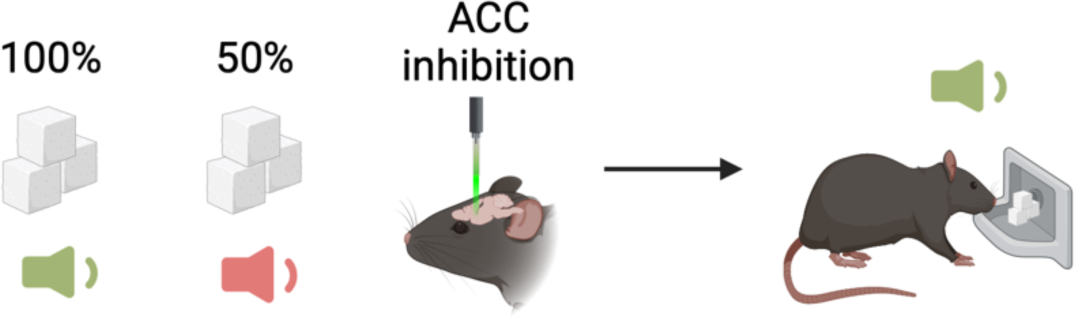

**Significance Statement:** We often make choices to gain information, even when the information cannot be used to change the outcome. Here we investigated the precise timing of the role of the anterior cingulate cortex (ACC) in decisions that involve seeking certain versus uncertain rewards. By optically inhibiting ACC neurons, we demonstrate that this region is crucial for maintaining persistence toward rewards signaled with certainty, without altering the motivational value of the reward itself. In a positive control experiment, we also confirm that ACC is important in effort-based choice. The findings reveal a common role for ACC in maintaining persistence toward certain and valuable rewards, necessary for making optimal decisions. These results have implications for understanding psychiatric disorders involving maladaptive reward-seeking behavior.

---

Individuals often are willing to pay a cost for information even when the information cannot be used to change the outcome. Prior research has shown that information-seeking behavior is modulated by a network involving frontocortical regions such as anterior cingulate cortex (ACC) and orbitofrontal cortex, along with subcortical regions like the striatum and pallidum (Bromberg-Martin & Monosov, 2020). Indeed, macaque ACC activity is one the earliest predictors of future choices suggesting it is important in establishing the drive to actively seek information from the environment (White et al., 2019). By comparison, research on the specific brain regions tracking information value in rodents is scarce (Bussell et al., 2023), as are inquiries about the causal mechanisms involved. Recently, we found that ACC was critical in information choice preference and that its mechanism was sex-dependent: previously rewarded information choices strongly predicted future choices in control animals, but not in female rats following ACC inhibition. These results revealed a causal role for ACC in decisions involving information (González et al., 2024).

To model information choices in nonhumans, several groups (Macias et al., 2021; Molet et al., 2012; Orduña, 2015; Pisklak et al., 2015; Stagner & Zentall, 2010; Vasconcelos et al., 2015) administer a task where rats choose between a more profitable option followed by a discrete cue, rewarded 50% of the time (*No information*) vs. a less profitable option followed by a cue that signals a certain outcome (*Information*). One of these cues (S+) always results in reward (100%) and the other (S-) is never rewarded (0%). In this procedure, the information follows the choice and does not affect the outcome and thus, any preference for that option suggests information is inherently rewarding (Cunningham & Shahan, 2019; Macias et al., 2021; Orduña, 2015; Stagner & Zentall, 2010). A preference for information could be due to an overweighting of the cue that signals certain reinforcement (S+) and/or an underweighting of the cue paired with the absence of reward (S-) (González & Blaisdell, 2021; González et al., 2023; McDevitt et al., 2016). However, behavioral manipulations involving either the S+ or S-have led to largely equivocal results, with more studies garnering support for overweighting the S+ (Fortes et al., 2016; Stagner et al., 2011).

To more precisely uncover the timing of the role of ACC in information choices, we studied decision making following optogenetic inhibition of this region during discrete trial epochs: On separate sessions during the presentation of the informative cue that fully predicts reward (S+), or during the intertrial interval (ITI). As a positive control for the effectiveness of optogenetic inhibition of ACC, we evaluated animals on effort-based choices involving high vs. low valued rewards, where its role has been documented extensively (Bailey et al., 2016; Floresco & Ghods-Sharifi, 2007; Hart et al., 2020; Hauber & Sommer, 2009; Mashhoori et al., 2018; Porter et al., 2019).

## Materials and Methods

### Animals

Twenty-four Long Evans rats (Rattus Norvegicus) *(N =* 14 females) were acquired from Envigo (Bioanalytical Systems, Inc., Indianapolis, IN). Two animals were not included in the analyses due to unilateral expression of the virus, one due to wrong placement of the optic fiber, and one on losing the headcap during the experiment (total N = 20, 10 females). A subset of animals (n=11) was also tested on a progressive-ratio (PR) task. Subjects were aged between post-natal day (PND) 90 and 120 at the start of the experiment. Subjects were paired-housed before and separated after surgeries, housed in transparent plastic tubs with wood shaving bedding in a vivarium maintained on a reverse 12-hr light cycle. Experiments were conducted at a minimum of six days per week during the dark portion of the light cycle. A progressive food restriction schedule was imposed prior to the beginning of the experiment to maintain rats at 85% of their initial free-feeding weights. Water was provided *ad libitum* in the homecage. The procedures used in this experiment were conducted under approval and following the guidelines established by the Chancellor’s Animal Research Committee at UCLA.

### Viral constructs

An adeno-associated virus AAV9 driving an inhibitory opsin archaerhodopsin T under the CaMKIIa promoter was expressed in neurons in ACC (n=15, AAV9-CaMKIIa-ArchT)-Green, packaged by Addgene, Addgene, viral prep #99039). A virus lacking the Arch-T gene and instead containing the fluorescent tag eYFP (n=5, AAV9-CaMKIIa-EYFP, packaged by Addgene) was infused into ACC as a null virus control. All animals underwent identical surgical procedures, and all behavior was conducted in a counterbalanced within-subject design with each animal serving as its own control between inactivation sessions. This allowed us to account for non-specific effects of surgery, exposure to AAV9, and non-specific effects of laser stimulation.

### Behavioral apparatus

Behavioral testing was conducted in Med Associates (East Fairfield, VT) operant conditioning chambers. Each chamber contained two retractable levers inserted to the left and right of a lower food delivery port in the front wall. A photobeam entry detector was positioned at the entry to the food port to detect head entries. A pellet dispenser connected to the food port delivered a single 45-mg sucrose pellet (Bio-Serv, Frenchtown, NJ). Tone, clicker and white noise generator were attached to individual speakers on the wall opposite the lever and magazine. A 3-watt, 24-volt house-light on the top of the back wall provided illumination. For optogenetic manipulations, the chambers were outfitted with an Intensity Division Fiberoptic Rotary Joint (Doric Lenses, Quebec, QC, Canada) connecting the output fiber optic patch cords to a 532nm Green laser (Shanghai Laser & Optics Century Co., Ltd. (SLOC), China) positioned outside of the chamber.

### Surgical procedures

After completing the training phase, rats were anesthetized with isoflurane for bilateral infusion of ACC with either an inhibitory opsin archaerhodopsin T (AAV9-CaMKII-ArchT)-Green (Addgene, Cambridge, MA, viral prep #99039) or eYFP (AAV9-CamKII-EYFP; Addene, Cambridge, MA). Craniotomies were created for the injection sites but also four extra holes were used to screw anchors. After the four screws were fixed in the skull, animals were infused with two bilateral sites of injections in ACC (0.3 µL at 0.1 µL/min AP: +3.7, ML: ±0.8, DV: -2.4 for a total volume of 0.3 µL per side). After infusion, the blunt needle was left in place for 10 min to allow diffusion. Using the same craniotomies, a stainless-steel fiber optic cannula ferrule (size: 1.25mm, 200 um Core, NA: 0.37, length: 2.5 mm; Thor Labs, Newton, NJ) was inserted in each side (AP: +3.7, ML: ± 1.0, DV: -2.1, inserted in a 15° angle). All measurements were taken from Bregma. Optic fibers were secured in place with cyanoacrylate, bone cement (C&B Metabond, Parkell Inc., Edgewood, NY) and dental acrylic (Patterson Dental, St. Paul, MN). Animals were provided with post-operative Carprofen (5 mg/kg, s.c.; Zoetis, Parsippany, NJ) daily for 5 d.

### Behavioral procedure

#### Pretraining

Before training, 10 sucrose pellets (0.5 g) were provided in the homecage to accommodate rats to the food rewards. On the first day, rats were trained to eat pellets from the pellet tray by delivering 1 pellet every 20 ± 15 s in the chamber for a total of 40 pellets. On days 2 and 3, rats were trained to lever press the left and right lever (lever presented in alternate order) in an autoshaping procedure. Reinforcements were delivered following each lever press. After pretraining, animals were either trained on the information choice task or the effort-based choice task, in counterbalanced order.

#### Information choice Training

The training stage of the task was comprised of two types of trials: choice and forced trials. In choice trials, rats were required to choose between two levers available simultaneously. A choice trial started with the simultaneous insertion of both levers and the houselight turning on. A choice was made by pressing a lever one time. After the choice was made, if the animal chose the Info option, both levers retracted and the houselight turned off. If the lever associated with the Info option was pressed, 20% of the time an auditory cue (S+) was presented for 30 s always ending with food; the other 80% of the time another auditory cue (S-) was on for 30 s never ending with food. If the rat chose the No-Info option, a third auditory cue (S3) was presented for 30 s and ended with food on half of the trials. On forced trials, only one lever was presented at a time, following the same contingencies described above. All trials were separated by a 10-s intertrial-interval (ITI) during which all stimuli were off. A single session constituted 54 trials, 18 choice trials, and 36 forced trials. The assignment of sounds to S+, S- or S3 and the lever side for each option was counterbalanced across animals. After 10 sessions, rats were connected to the patch cords without any stimulation to habituate them to move around and perform the task with the optical tether (200 µm, 0.22 NA, Doric Lenses) for another 6 sessions, with no light delivery. Thus, training was a total of 16 sessions.

#### Information choice test

During two consecutive sessions, rats were tested on the task as described above but this time receiving optogenetic inhibition during either the 30s S+ presentation or on 10% of ITI periods (to equate to the duration of other trial epochs and minimize potential damage due to prolong photoinhibition). Green light was delivered into ACC, which inhibited pyramidal neurons in this region during these trial epochs in rats expressing ArchT and not eYFP controls. During each presentation, a green light (532 nm; 10 mW) was delivered via a laser (532nm; Shanghai Laser and Optics Century Co., Shanghai, China) connected through a ceramic mating sleeve (2.5 mm; Precision Fiber Products, Chula Vista, California) to the ferrule implanted in the rat. Optogenetic parameters were chosen considering previous work on the duration of continuous inhibition in ArchT-expressing neurons (up to 1-15 min) (El-Gaby et al., 2016; Huff et al., 2013) compared to our 30-sec inhibition, and considering that cell body inhibition as we conduct here is more selective, limiting off-target effects compared to terminal inhibition (Lafferty & Britt, 2020). Light leakage from laser output was prevented using 5-cm long black shrinktube shielding over the connected patch cord and cannula ferrules. Between each condition, animals received a baseline session, in which no inhibition was administered but animals were connected to the optical tether as before. The order of inhibition during the three different epochs was counterbalanced (**Figure 1B**).

**Figure 1.**
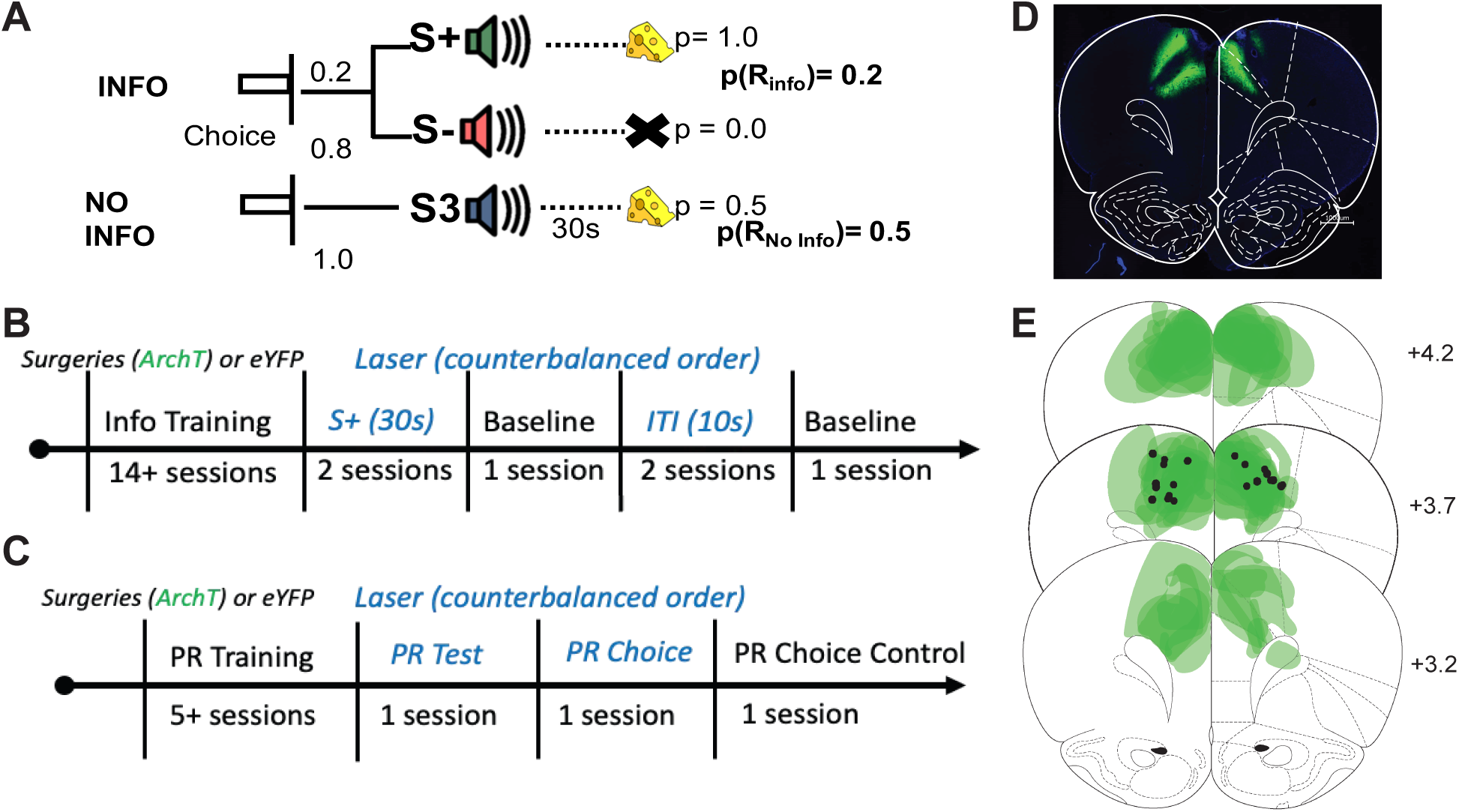
Information choice task, Effort-based choice task, timelines, and histology. **(A) Info Choice task.** Rats choose between two levers (left or right). If the rat chooses the ‘Info’ option, on 20% of the trials, tone S+ plays for 30s terminating in the delivery of one sugar pellet; the other 80% of trials tone S-plays, terminating in no food. If the rat chooses the ‘No-Info’ option, a third tone S3 plays for 30s, terminating in one sugar pellet on 50% of the trials. A session consisted on 54 trials evenly split between choice, forced-info and forced no-info trials. Only one lever was available on forced trials. **(B) Timeline Info task.** Rats underwent surgery for virus and fiber implant prior to any training. Conditions with optogenetic inhibition were during the S+ and ITI, counterbalanced and alternated with a no-laser session (baseline) between conditions. During the S+ condition, ACC inhibition occurred in choice and forced-info trials, during the duration of the cue S+ (30 s). During the ITI condition, ACC inhibition occurred on 10% of the 10-s ITI epochs. **(C) Timeline PR task.** Rats underwent surgery for virus and fiber implant prior to any training. During PR Test and PR Choice, ACC inhibition occurred during sucrose pellet retrieval. PR Test was just as the training session. In PR Choice and PR Choice Control, animals were presented with a ramekin of 20 g of chow. The order of testing was counterbalanced across animals. A subset of animals experienced both tasks, in counterbalanced order. **(D) Photomicrograph.** Representative placement of inhibitory opsin ArchT (green) and fiber placement at Anterior-Posterior (AP) level +3.7 for ACC relative to Bregma. **(E) Reconstructions.** Reconstructed placements of inhibitory opsin ArchT virus (green) and optic fiber placement (black) across all animals, for AP level +4.2, +3.7 and +3.2 relative to Bregma, targeting ACC.

#### Progressive-ratio (PR) Training (for Effort-based choice)

Animals were trained on a progressive ratio (PR) schedule where the required number of presses for each pellet increased according to the formula:

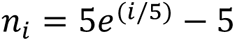

As described inHart et al. (2020), the effort requirement increased for the rat on earning 5 pellets where 𝑛_𝑖_ is equal to the number of presses required on the i^th^ ratio, rounded to the nearest whole number after 5 successive schedule completions. Each session started with the insertion of the left lever. The completion of a ratio requirement was marked by a light, which remained on until the animal collected the pellet from the food tray. A session lasted 90 min. The training lasted a minimum of 5 sessions and continued until stability was reached, defined as the average number of lever presses in a session not varying by more than 15% across the last two sessions.

#### PR Test

Animals experienced two test sessions in which ACC was inhibited, using the same parameters described in the Info choice test. In one test, animals underwent a PR session similar to the training phase, during which inhibition occurred during pellet retrieval, and where sucrose pellets were the only option. Our previous imaging of single cells in rat ACC revealed a phasic increase in activity during reward collection of the preferred reward(Hart et al., 2020), thus, we predicted that inhibiting ACC during this trial epoch would be particularly effective in changing effort-based decision making. In another test session, animals received a PR Choice session where the task occurred as described, but rats also had free access to 20 g of chow in a ceramic ramekin positioned on the right side of the food port (opposite the lever location, along the same wall as the pellet tray). For both tests, the onset of the inhibition occurred at pellet delivery and lasted for 2 seconds or until the animal collected the pellet or pressed the lever (whichever occurred first). Animals also underwent a PR Choice control session identical to PR Choice but without laser inhibition. The order of test sessions was counterbalanced (**Figure 1C**).

### Histology

At the conclusion of the experiment, rats were euthanized with an overdose of sodium pentobarbital (Euthasol, 0.8 mL, i.p.; VetOne, Paris, France) and transcardially perfused with phosphate buffered saline (PBS) followed by 10% buffered formalin acetate. Brains were extracted and post-fixed in this solution for 24 hours followed by 30% sucrose immersion.

Tissue was sectioned in 40-µM thick slices and cover slipped with DAPI mounting medium (Prolong gold, Invitrogen, Carlsbad, CA) visualized using a BZ-X710 microscope (Keyence, Itasca, IL), and analyzed with BZ-X Viewer software. A representative image of the viral expression and implant tracks in the Arch-T group is shown in **Figure 1D**. Reconstructions of the viral expression across all Arch-T expressing animals are shown in green with black dots indicating the location of the bottom of the optic fiber implants across animals (**Figure 1E**).

### Data Analysis

All data processing and visualization were conducted using R Studio and GraphPad Prism 10 (GraphPad Software LLC, San Diego, CA). All data were subsequently analyzed via custom-written code in MATLAB (MathWorks, Inc., Natick, MA). For the Info choice task, we analyzed three main variables: 1) preference for the Info option (Info choice divided by the total number of choices), 2) latency to choose (the time from the onset of the trial until lever press), and 3) port entry rate (the number of entries into the food port) during the 30-s cue duration. For all analyses, we used mixed-effects General Linear Models (GLM) (*fitglme* function; Statistics and Machine Learning Toolbox) to more rigorously handle data interdependencies (de Melo et al., 2022; Yu et al., 2022) such as cue exposure that depended on individual rat’s choice. Preference data were analyzed first in omnibus GLM that included all factors (sex, trial epoch, laser on/off) and all groups (virus: Arch-T vs. control). Sex was included as a moderator in training and a covariate at test. Follow-up analyses were further pursued pending significant interactions in the full model with planned comparisons: S+, S-, S3 and PR, PR Choice. Coding of variables for GLMs was as follows: 0 = females and 1 = males; 0 = control and 1 = ArchT; 0 = ITI, 1 = S+; 0= laser off (no laser), 1= laser on (laser on in the current trial). Individual rat was entered as a random factor. Latency analysis included the described variables in addition to the within-subject variable: trial-type (forced-info, forced-no info and choice). Port entry rate analysis included cue as a crossed, within-subject variable (S+, S- and S3).

For the PR task we analyzed three main measures: 1) Mean lever presses in a session, 2) Breakpoint (BR), that is, highest incomplete PR ratio, and 3) Latency to collect reward. As in the information task, we used GLM planned comparisons to probe significant interactions, when observed. To assess correlations between information preference and performance on the effort task we conducted Spearman rank analyses. For this we included only animals that had been involved in both tasks on only laser-inhibited trials, and included both types of effort contexts (PR and PR Choice). All code and data can be found in the following repositories: https://github.com/izquierdolab and https://gin.g-node.org/aizquie.

## Results

### Information choice training: Rats acquire a stable preference phenotype, and respond faster in choice trials

*Preference.* We found that rats exhibit a stable preference phenotype that does not change by the end of training. Training data were analyzed using preference across sessions. A GLM was performed for the outcome variable, information (*Info*) preference, using session as within-subject factor, sex as a between-subject factor, and individual rat as a random factor using the formula: *γ ∼ [1 + sex*session + (1 + session| rat)].* We found no significant predictors or interactions with session [GLM: β_session_ = 0.005, *t*(290) = 0.65, *p* = 0.51], sex [GLM: β_sex_ = 0.025, *t*(290) = 0.40, *p* = 0.68], or session*sex [GLM: β_session*sex_ = 0.0002, *t*(290) = 0.02, *p* = 0.98].

The absence of a significant effect of session may be due to the high variability of choices in the development of a preference. To address this, we classified animals based on the phenotype exhibited during the last three sessions of training: Info preferring (preference above 0.6), No Information preferring (*No Info*) (preference below 0.4), and Indifferent (between 0.4 and 0.6). Using phenotype as a fixed factor moderator in the formula *γ ∼ [1 + sex*phenotype*session + (1 + session| rat)],* we found a significant effect of session [GLM: β_session_ = 0.033, *t*(286) = 2.71, *p* = 0.007] and a significant session*phenotype interaction [GLM: β_session*phenotype_ = -0.023, *t*(286) = - 2.68, *p* = 0.007]. This interaction indicates that preference changed differentially across sessions of training. Animals began their preference around the indifference point 0.5 but then diverged, with a group of animals remaining at the indifference point and the other two groups developing a strong preference towards/against the Info option (**Figure 2A**). When selecting the last three sessions of training to check for stability, use of the same formula led to no significant predictors or interactions, confirming animals’ preference was stable by the end of training.

**Figure 2.**
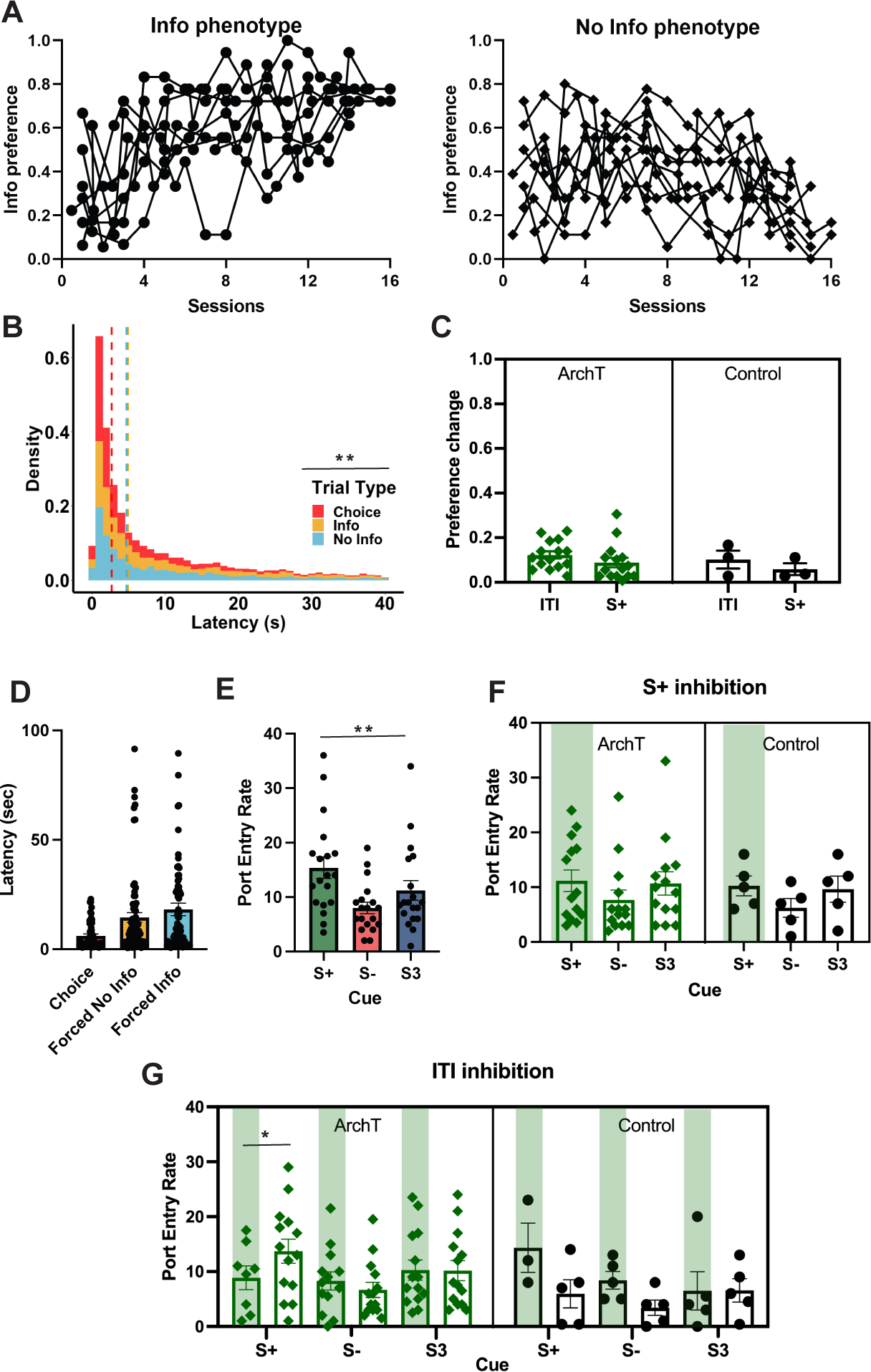
Optogenetic inhibition of ACC prospectively reduces reward expectation during the Info (S+) cue. **(A)** Establishment of phenotype in animals during training. Each line represents an individual rat. **(B)** Density of the total number of observations for latency to choose by trial type collapsed across conditions during training. Dashed lines refer to median latency. Median latency to choose in choice trials (red line) was faster than in forced trials (gray line). **(C)** Mean absolute preference change from baseline (no laser) during inhibition of S+ cue or ITI presentation, with individual rats represented as scatter. **(D)** Median latency to choose and during forced No Info and forced Info trials at test. **(E)** Median port entries during the 30 s cue presentations (S-, S+ and S3) for all animals. Individual circles correspond to median responses for each animal. **(F)** S+ inhibition: Median port entries during each cue presentation on laser on (green vertical stripes) and off (no stripes) for ArchT (left, green) and control (right, black) animals, with individual rats represented as scatter. **(G)** ITI inhibition: Median port entries during each cue presentation following laser on (green vertical stripes) and off (no stripes) for ArchT (left, green) and control (right, black) animals, with individual rats represented as scatter. A decrement in port entries during S+ was observed following inhibition in the ArchT group, but not the Control group. There was a general increase in port entry rate in laser on trials in the Control group, but it was not specific to a cue. Error bars denote ± SEM. **p<0.01, *p<0.05. ArchT n=11, Control n=5.

#### Latency to choose

We found that rats are faster to make choices than to complete forced trials, corroborating prior research (González et al., 2023; González et al., 2024). We conducted a GLM on median latencies using trial type (forced Info, forced NoInfo, and Choice) as within-subject factors, sex as between-subject factor, and individual data as a random factor, formula: *γ ∼ [1 + trial type*sex + (1 + trial type| rat)]*. The results yielded a significant main predictor of trial type [GLM: β_trial type_ = 5.19, t(68) = 3.79, *p* < .001], but no effect of sex [GLM: β_sex_ = 1.43, t(68) = 0.52, *p* = 0.60], or interaction [GLM: β_trial type*sex_ = -.167, t(68) = -0.60, *p* = 0.55]. Indeed, rats are faster during choice trials (Median_Choice_= 4.4 s, SEM ± 0.81) than forced trials (Median_Forced Info_= 11.4 s, SEM ± 2.46; Median_Forced NoInfo_= 7.6 s, SEM ± 2.64) (Figure 2B).

#### Port entry Rate

Port entry rate is an indication of differential reward expectation for S+ vs. S-. We analyzed the median port entry rate (entrance to the feeder port) for each 30-s cue duration for the last three sessions of training. A GLM was conducted for median port entries using sex as a between-subject factor, and individual rat crossed with cue (S+, S- and S3) as a random factor, according to the following formula: *γ ∼ [1 + sex*cue + (1|rat:cue)]*. Cue was included in this way to account for the fact that rats experienced each cue a different number of times given the inherent contingencies of the task: S+ was presented on only 20% of all Info trials but it also depended on the rat’s choices (for instance, a rat that chooses the No Info option exclusively would have an overrepresentation of the S3 cue and very few S+ trials). We found no effect of cue or sex on port entry rate (p > 0.64). We have previously reported the greatest responses for the S+ and lowest responses to S- (González et al., 2024). However, in that study, the cue duration was 60s, affording a longer window for differences to emerge.

### Information choice test: Optogenetic inhibition of ACC neurons decreases port entry rate during S+ cue, but does not affect overall information preference or latencies

*Preference*. Info preference during test sessions was analyzed for both epochs. Mean values were used for S+ and ITI inhibition sessions. The GLM formula was *γ ∼ [1 + virus*laser*epoch+sex+ (1 + epoch+laser| rat)]*, with epoch (S+ or ITI inhibition) and laser entered as within-subject factors, and sex as a covariate. None of these factors or their interactions significantly predicted Info preference. Given that we observed high variability in the initial preference, any detectable changes in preference depended on each rat’s baseline. Therefore, we calculated the absolute change in preference during each optogenetic inhibition session minus the preference during baseline, identical to the analysis in González et al. (2024). The same GLM formula was used but instead we used the *absolute preference change* as the outcome of interest. No significant predictors or interactions between factors was found.

Similarly, these same factors did not predict *phenotype*. In summary, optogenetic inhibition of ACC did not affect overall preference or change in preference phenotype (**Figure 2C**).

#### Latencies

Optogenetic inhibition of ACC could affect performance measures such as latencies, not just preference *per se*. To test this, we analyzed median latencies to choose during test sessions. Here we used the formula: *γ ∼ [1 + virus*epoch*trial type*laser +sex+ (1 + epoch + trial type+ laser| rat)]*; where virus and sex were between-subject factors, epoch (S+, ITI, and Baseline), trial type (Choice, forced Info and forced No info), and laser (laser on, off) were within-subject factors, and individual rat was a random factor. The results revealed no significant predictors or interactions (**Figure 2D**).

#### Port entry rate

Median reward port entries were analyzed for S+ and ITI epochs using the following GLM formula: *γ ∼ [1 + virus *cue*laser + (1 + laser| rat:cue)]*, and separately for the baseline condition with laser and virus not included in the formula because laser was not yet introduced: *γ ∼ [1 + cue + (1 | rat)].* The analysis of baseline sessions across all animals resulted in a significant effect of cue [GLM: β_cue_ = -2.09, *t*(55) = *p* = .005] (**Figure 2E**). The analysis of S+ inhibition (with cue left out of the model) resulted in no significant effects (*p* > .17) or interactions (*p* > 0.88) (**Figure 2E**). However, inhibition during the ITI epoch resulted in a significant effect of laser [GLM: β_laser_ = 9.87, *t*(97) = 2.66, *p* = .009], a virus*laser interaction [GLM: β_virus *laser_ = -14.97, *t*(97) = -3.45, *p* = .0008], and virus*cue*laser interaction [GLM: β_virus*cue*laser_ = 5.21, *t*(97) = 2.72, *p* = .007]. Due to these interactions, we separately investigated the effect of inhibition in the ITI epoch by virus, where we used this GLM formula for predicting median port entry rate: *γ ∼ [1 + cue*laser + (1 + laser| rat:cue)]*.

The analysis for the ArchT, but not controls, resulted in a laser*cue interaction [GLM: β_laser*cue_ = 2.13, *t*(73) = 2.15, *p* = .035], indicating the inhibition differentially affected responsivity to cues. Planned comparisons for each cue resulted in a significantly lower port entry rate during the subsequent S+ [GLM: β_laser_ = -4.65, *t*(20) = -2.33, *p* = .03], but not S- (*p* = .20) or S3 (*p*= .89) (**Figure 2F**). The analysis of the control virus group resulted in a significant effect of laser [GLM: β_laser_ = 9.119, *t*(24) = 2.48, *p* = .020], with animals checking the food port more (not less) following all “laser on” trials, irrespective of the cue presented. This effect was not explained by animals being distracted by residual light, given that the responses during cue presentation occurred after the laser was experienced and could instead reflect enhanced arousal and activity levels following laser stimulation in controls. In sum, ACC inhibition led to a prospective decrease in reward checking specifically during the S+, indicating that animals did not exhibit the appropriate expectation of a certain reward.

### Progressive Ratio: ACC inhibition reduced effort and decreased breakpoint, without affecting latency to retrieve reward

*Total lever presses*. We analyzed total lever presses during sessions with laser on during the reward collection period. This epoch was selected based on previous work from our lab implicating phasic increases in population activity in ACC for higher-valued rewards as essential for effort-based decision making (Hart et al., 2020). In these analyses, virus was a between-subject factor, test (PR Test and PR Choice Test) was a within-subject factor, sex was a covariate, and individual rat was entered a random factor. The GLM formula was as follows: *γ ∼ [1 + virus* test + sex + (1 + test| rat)].* We found a main effect of test [GLM: β_test_ = -1229.6, *t*(17) = -4.02, *p* = .0008], an effect of virus [GLM: β_virus_ = -2191, *t*(17) = -2.42, *p* = .027] but no effect of sex (*p* = .30) nor interactions (*p*= .19), indicating that responses during the PR Test were higher than during PR Choice Test, but also that overall responses were lower for the ArchT virus group compared with the control animals (**Figure 3A**). We also compared performance during sessions with no laser (last session of training and PR Control Choice). For this, the GLM formula was identical to the previous analysis, and here we found a significant effect of test [GLM: β_test_ = -678.4, *t*(17) = -4.86, *p* = .0001], also resulting in a higher level of responding during PR training than PR Control Choice, but no effect of virus (*p* = .09), sex (*p* = .22), or interactions (*p* = .28), **Figure 3B**. These results collectively show the expected effect that animals tend to work more vigorously when sucrose pellets (i.e., in PR training or PR Test, but not PR Choice or PR Control Choice) are the only option. More importantly, these data indicate that ACC inhibition reduced effort, and that this decrement in effort was not due to a general reduction in motivation *per se*, given that rats still lever press more when sucrose was the only reward (i.e., the decrement was proportional to their initial performance in the given test). If ACC inhibition was affecting motivation, we would expect a similar level decrement in PR Test and PR Choice.

**Figure 3.**
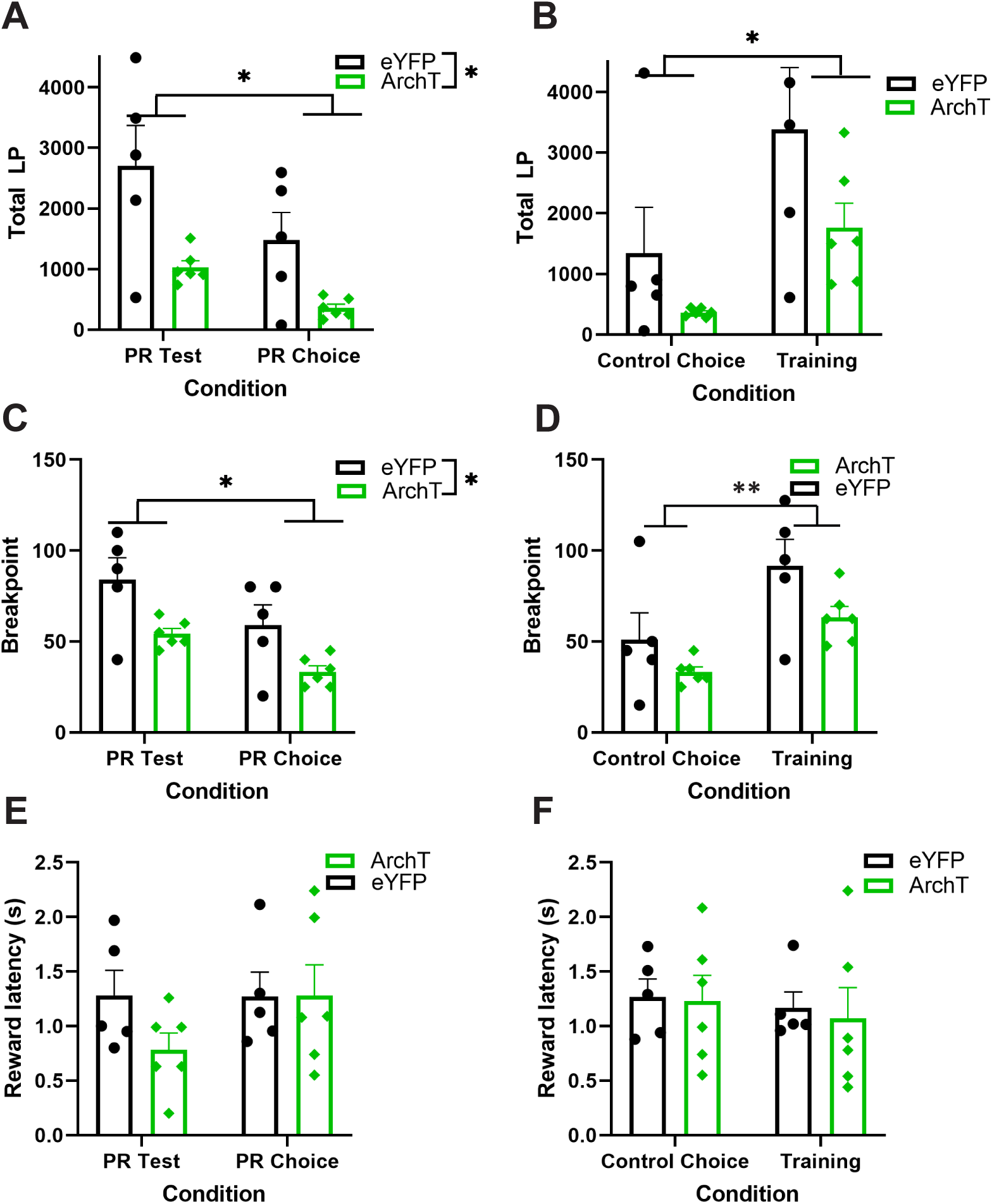
Optogenetic inhibition of ACC during PR and PR Choice decreases effort without affecting motivation. **(A)** Mean total lever presses during PR test and PR choice for ArchT (green) and control (black) animals. Individual rats represented as scatter. **(B)** Mean total lever presses during control Choice (PR choice with laser off) and last session of training for ArchT (green) and control (black) animals. Individual rats are represented as scatter. Lever presses during Control choice test were lower than during training, with no difference between virus group. **(C)** Mean BP during PR test and PR choice for ArchT (green) and control (black) animals. Responses were lower during PR choice than PR test, but also lower for ArchT animals. **(D)** Mean BP during Control choice and Training (no inhibition) for ArchT (green) and control (black) animals. Individual rats are represented as scatter. A significant difference of test but not virus was found. **(E-F)** Mean latency to collect reward during inhibited trials (PR Test, PR Choice) and during noninhibited trials (control choice and training) for ArchT (green) and control (black) animals, with individual dots corresponding to median latency for each rat. No difference in reward latencies were observed. *p<0.05, ***p<0.01. ArchT n=6, Control n=5.

#### Breakpoint

The last incomplete PR step was noted as the breakpoint. We analyzed breakpoint for the two tests with inhibition (PR Test and PR Choice), with an identical GLM formula as for lever presses: *γ ∼ [1 + virus* test + sex + (1 + test| rat)].* This analysis also yielded a main effect of virus [GLM: β_virus_ = -33.30, *t*(17) = -2.32, *p* = .033], and test [GLM: β_test_ = -25, *t*(17) = -4.92, *p* = .0001], but no effect of sex (*p* = .46) nor interactions (*p* = .55), **Figure 3C**. For the test sessions without laser (PR Training and PR Control Choice) we used the same formula as above. Here we found an effect of test [GLM: β_test_ = -13.5, *t*(17) = -7.04, *p* = 2.00^e-06^], but no other effects (*p* > .05) or interactions (*p* = .18), **Figure 3D**. Taken together, this confirms that the decrement in effort following ACC inhibition depended on the availability of another source of reward.

#### Latencies

We analyzed median latencies to collect the reward. As before, we analyzed the two tests where inhibition occurred using the following GLM formula: *γ ∼ [1 + virus* test + sex + (1 + test| rat)]*, where *γ* was median latency to collect reward, virus was a between-subject factor, test was a within-subject factor, sex was entered as covariate, and individual rat was entered as a random factor. Comparing PR Test and PR Choice Test, we found a significant effect of sex [GLM: β_sex_ = -0.40, *t*(17) = -2.45, *p* = .026], where females were faster at collecting reward than males, yet no other significant predictors (*p* > .089) or interactions (*p* = .22) were observed, **Figure 3E**. Similarly, for PR training and PR Control Choice no significant effects (*p* > .13) or interactions were found (*p* = .74), **Figure 3F**. Collectively, these results suggest that the decrement in effort cannot be explained by a general reduction in motivation to the rewards, where we would expect an increase in latencies to collect pellets.

### Correlations between Info Preference and Effort

Spearman rank correlation coefficients were computed to assess the relationship between the main measure of information preference and all other main dependent measures in the PR task (both PR and PR choice). Expectedly, we found significant correlations within the effort-based task itself, with positive correlations between median lever presses and total lever presses (r = 0.899, *p* = 0.028), and total lever presses with breakpoint (r = 0.986, *p* = 0.006). Interestingly, the only significant association with information preference was median reward latency in the PR task (r = -0.899, *p* = 0.028), which was a negative association: animals that had a strong information preference were quicker to collect reward on the PR task.

## Discussion

We investigated the role of ACC in information and effort choices. Previous research has highlighted the involvement of ACC in decision-making processes, particularly in evaluating effort costs (Bailey et al., 2016; Cai & Padoa-Schioppa, 2021; Fatahi et al., 2020; Hart et al., 2020; Hart et al., 2017; Hosokawa et al., 2013), so we used this behavior as a positive control for the efficacy of our optogenetic manipulation in Info choices. In previous studies, we and others have found ACC to be important in maintaining a representation of the value of *both* options in a 2-alternative choice paradigm (Hart et al., 2020; Hart et al., 2017; Mashhoori et al., 2018). The present results from both tasks suggest that ACC is required to maintain the appropriate reward expectation, either for the higher-valued reward of sucrose (in the effort task), or for the more certain reward (in the Info task). Interestingly, inhibiting ACC neurons decreases this stay-on-goal behavior without changing motivational value *per se* because reward latencies were unaffected by inhibition. Based on the results of this experiment, we suggest that this may be the mechanism by which stable preference is maintained in Info choices (González et al., 2024). The ITI would be the crucial epoch in the task where previous trial experience gets consolidated to update the course of future action. Indeed, a preference for information on the Info task was correlated with shorter reward latencies in the effort task, indicating these two measures are related via reward expectation.

Despite the effect of inhibition on reward expectation at the trial level, we were surprised to find no change in global Info preference. However, in order to equate overall inhibition time between S+ and ITI epochs, we only inhibited ACC on 10% of the 10-s ITIs. Had we inhibited ACC neurons on more of these ITIs, we may have observed an effect on overall preference. In keeping with the literature, we did observe distinct behavioral phenotypes for information choices: most rats either exhibited a strong preference for the Info option or preferred the No-info option. Almost no animal remained indifferent to the choice options. This divergence suggests that information may be valuable, but when there is a cost, such as via a potential loss of reward opportunities, there are individual differences in that cost sensitivity.

Future experiments could modulate or titrate this cost sensitivity as a continuous variable. Our present findings may intersect with the literature on sign- vs. goal-tracking as there are relevant discoveries worth noting in that vein. For example, 1) both uncertainty reduction and conditioned reinforcement influence information choice (Ajuwon et al., 2023); 2) strong stimulus-reward predictive utility promotes information preference (Chow et al., 2017); and 3) individual differences in sign- vs. goal-tracking do not fully predict information preference in rats (Lopez et al., 2018). We expected that ACC would be robustly involved in this task due more to its involvement in uncertainty monitoring rather than any role in conditioned reinforcement. Unlike measures of decision confidence assessed by temporal wagering for reward (Lak et al., 2014; Stolyarova et al., 2019), in this task the rat makes a choice prior to cue presentation and the auditory cue remains on from the moment the rat has selected the option until the outcome is revealed (i.e., reward or no reward). As a result, animals are not relying on memory and do not need to keep the cue ‘in-mind.’ Therefore, because inhibition of ACC neurons reduces subsequent checking of the reward port when there is certain to be a reward (S+), this is evidence neurons in this region support reward expectancy rather than uncertainty monitoring in this task. Better goal maintenance could facilitate exploration, grooming, or resting etc., ultimately supporting more behavioral flexibility under certainty.

We and others (Cox et al., 2023) have previously proposed that ACC is important in accessing the relative value of each option, which as mentioned above requires a representation of both outcomes. In support of this, we find that ACC inhibition decreased reward expectation for a certain reward by affecting goal maintenance. Recent work by Vasquez et al. (2024) proposed that ACC is important for attention-related processes, contributing to the ability to “stay on task” (Vázquez et al., 2024). Recent work by Regalado et al. (2024) is also consistent with our findings. This group investigated the role of neural activity in ACC when mice self-initiate trials to learn cue-reward contingencies. Authors found that neural activity in orbitofrontal cortex-to-ACC projections increased progressively with consecutive unrewarded cues and peaked at a reward-associated cue, keeping the animal “on track” in the task. Most relevant to the present study is that ACC maintains the relevant goal information to allow for such goal-directed behavior.

On the effort choice task, ACC inhibition had no impact on the speed of reward collection, but did reduce effort. This aligns with the extensive literature implicating ACC in effort-related decision-making, supporting the idea that ACC activity is integral to sustaining high-effort behavior for high-value rewards (Azab & Hayden, 2017; Bailey et al., 2016; Cai & Padoa-Schioppa, 2021; Fatahi et al., 2020; Hosking et al., 2014). Consistent with what was observed in the Info choice task, the lack of changes in latencies to collect reward suggest that the motivational value of the reinforcer remained intact. Furthermore, the decrement in effort during ACC inhibition was proportional to the type of test, that is, the effort was lower when an alternative source of reward was introduced (PR Choice) compared to when only sugar pellets were available (PR Test). If ACC inhibition was involved in altering motivation, a similar level of effort would have been observed if either one or two sources of reward were present. Instead, the pattern of responses was preserved and did not affect speed of retrieval, suggesting a disruption in maintaining the goal of the better option (i.e., sucrose). We have previously shown sucrose is indeed the better option when the choice is between chow and sucrose pellets (Hart et al., 2020; Hart et al., 2017).

In summary, while ACC inhibition did not directly alter Info preference or reward latencies, it significantly affected reward expectation during an informative (S+) cue and attenuated effort-based choice. These results underscore a nuanced role for ACC in information-seeking behavior and effort-based choice via a common stay-on-goal mechanism. Future studies should explore more precise temporal and spatial dynamics of ACC activity to further elucidate its role in these complex behaviors.

**Table 1.** Info Preference and Effort-based Choice (Spearman Rank Correlation Coefficients)

|  | InfoPref | Median<br>LP | Total<br>LP | Reward<br>Latency | Breakpoint |
| --- | --- | --- | --- | --- | --- |
| InfoPref | 1.0 |  |  |  |  |
| Median LP | 0.388 | 1.0 |  |  |  |
| Total LP | 0.429 | **0.899\*** | 1.0 |  |  |
| Reward Latency | **-0.899\*** | -0.662 | -0.580 | 1.0 |  |
| Breakpoint | 0.377 | **0.956\*** | **0.986\*\*** | -0.588 | 1.0 |

\*p<0.05, \*\*p<0.01

## Acknowledgements

We thank members of the Izquierdo, Sharpe, and Wassum labs for early feedback and suggestions on these experiments.

Authors report no conflict of interest

